# Determination of Optimal Nutrient Levels in Baru Seedlings (Dipterix alata Vog.) Using the Mathematical Chance Method

**DOI:** 10.1101/2023.10.20.563291

**Authors:** Daniel Kusano, Leia Larson, Marcos Antonio Camacho

**Affiliations:** SENAR; PREFEITURA DE AQUIDAUANA; UEMS

## Abstract

This work aimed to determine the optimal ranges by the Mathematical Chance (ChM) method in baru seedlings. To determine optimal ranges by ChM, the nutritional information from an experiment conducted in a protected environment from October 13, 2011, to February 11, 2012, in Selvíria-MS and for seedling production, used a nursery with 50% light reduction polypropylene mesh and 15 x 21.5 cm plastic bags, which have a capacity of 1.8 L, which were filled with different substrate combinations. The ranges found for macronutrients with the ChM method were, in g kg-1, from 19.06 to 28.14; 1.52 to 3.38; 17.9 to 24.0; 1.76 to 3.54; 0.74 to 1.50; 1.26 to 2.26 for N, P, K, Ca, Mg and S, respectively. The ranges found for micronutrients with the ChM method were, in mg kg 1, from 21.79 to 33.59; 1.0 to 21.40; 75 to 104.4; 14.0 to 82.8; 18.0 to 24.4 for B, Cu, Fe, Mn, and Zn, respectively.

## Introduction

Baru (Dipterix alata Vog.) is a native species of the Brazilian Cerrado (AJALLA et al., 2012) and offers multiple utilization potential, including food, wood, and land restoration (MARTIMOTTO et al., 2012). Despite the potential use of this species and many other native Cerrado species, research is still in its early stages, and there are several knowledge gaps to be filled.

However, information about this species is limited, with some studies evaluating seedling production (AJALLA et al., 2012; SANTOS et al., 2019) and nutrient absorption (SILVA et al., 2016; BONI et al., 2016; SOUZA et al., 2018). Little is known about the nutritional requirements of this species (CARLOS et al., 2014).

To enhance the nutritional study of plant species, it suggested that leaf analysis should be more widely used in fertilizer recommendation programs (WADT & SILVA, 2016). Techniques such as leaf analysis are essential to identify nutritional deficiencies, especially for crops without well-established references, and data interpretation from various methods (CAMACHO et al., 2012b).

The mathematical chance method (ChM) determines sufficiency ranges among the established nutritional diagnosis methods. To achieve maximum productivity, it establishes critical levels, optimal values, and sufficiency ranges based on nutritional monitoring data (WADT et al., 1998).”

The ChM classifies the leaf nutrient concentrations in ascending order with the productivity obtained in the respective plots where the samples are. After a series of calculations, it estimates the range of nutrient content where a higher probability of productivity response is expected (URANO et al., 2006).

In this context, this study aimed to determine the normal nutrient ranges for baru seedlings using the mathematical chance method based on data from a limited population in which the variation factor was the substrate composition.

## Materials and Methods

For the generation of optimal ranges using the ChM, nutritional information was derived from an experiment conducted in a controlled environment from October 13, 2011, to February 11, 2012, in the municipality of Selvíria-MS, located at latitude 20°25’ S and longitude 51°21’ W.

The region’s climate is classified as Aw according to Köppen, characterized as hot tropical with summer rains. The average annual temperature is 24.5 °C, and the average annual precipitation is 1,232 mm.

A nursery with polypropylene screens with 50% light reduction and plastic bags measuring 15 x 21.5 cm, with a capacity of 1.8 L, was used to produce the seedlings. These bags were filled with different substrate combinations, as described in Larson et al. (2018). Soil samples from the 0-20 cm clayey layer to moderately clayey typical dystrophic Red Latosol, hypodystrophic, aluminum-rich, kaolinitic, ferruginous, deep, and moderately acidic.

The plants were sampled 120 days after emergence by collecting the third fully expanded leaf of the baru seedlings to establish reference parameters for nutritional analysis. The evaluated nutrients in the samples included N, P, K, Ca, Mg, S, B, Cu, Fe, Mn, and Zn. The maximum and minimum values and the coefficient of variation for each nutrient are in Table 1. The productive parameter adopted was the dry mass of the aerial part (MSPA) of baru seedlings. To adjust the method, we divided the population into two classes: MSPA production above 3.84 g per plant, defined as the high MSPA production subpopulation, representing 80% of the maximum MSPA production, and the other below this.

**Table 1.** Minimum and maximum values found in the database, coefficient of variation (C.V.), class width, number of classes, and standard deviation for each evaluated nutrient.

| Nutrient | Minimum | Máximo | Class Width | Number of Classes | C.V. |
| --- | --- | --- | --- | --- | --- |
|  |  | g kg<sup>-1</sup> |  |  | (%) |
| N | 16.03 | 28.84 | 2.56 | 5 | 19.47 |
| P | 1.52 | 4.61 | 0.62 | 5 | 35.84 |
| K | 8.5 | 24 | 3.1 | 5 | 36.49 |
| Ca | 1.76 | 10.68 | 1.78 | 5 | 57.52 |
| Mg | 0.74 | 2.66 | 0.38 | 5 | 36.35 |
| S | 1.26 | 3.75 | 0.5 | 5 | 35.75 |
|  |  | mg kg<sup>-1</sup> |  |  |  |
| B | 21.79 | 51.28 | 5.9 | 5 | 18.03 |
| Cu | 1 | 52 | 10.2 | 5 | 116.98 |
| Fe | 75 | 222 | 29.4 | 5 | 30.59 |
| Mn | 14 | 358 | 68.8 | 5 | 69.04 |
| Zn | 18 | 34 | 3.2 | 5 | 16.43 |

For each nutrient under study, leaf concentrations were classified in ascending order and divided into some classes defined as the square root of the number of observations. The value intervals for each class were then determined by dividing the range of nutrient concentrations in question by the number of established classes, following the approach of Wadt et al. (1998). In each class, calculate the mathematical chance using the equation:

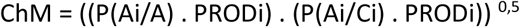

In which:

ChM = mathematical chance in class “i”

P(Ai/A) = the frequency of high productivity plots in class “i” relative to the overall total of high productivity plots (A = ΣAi)

P(Ai/Ci) = the frequency of high productivity plots in class “i” relative to the total number of plots in class “i”

PRODi = the average productivity of high productivity plots in class “i”

The number of classes was defined based on the different class ranges for each nutrient. The distribution of classes is in Table 1.

After calculating the ChM for each class, it determined the optimal range for each nutrient corresponding to the classes of values with the highest mathematical chance values. The lower limit was considered the critical level, and its median value was the optimal nutrient content (URANO et al., 2007).

## Results and Discussion

The highest ChM value found for nitrogen (N) was 4.14 g per plant, corresponding to the optimal range (WADT et al., 1998). However, by adopting only the optimal range as the sufficiency range, the recommendation would be in the range of 25.12 to 28.14 g of N per kg of MSPA, which is a very narrow range and may not precisely correspond to the crop’s sufficiency range (Table 2). Grouped as the ChM values, values above 2.05 g per kg were considered when determining the range. Therefore, for N, the ChM values were determined between classes 2 and 4 (19.06 to 28.14 g per kg), as proposed by Camacho et al. (2012a). Boni et al. (2016) reported N content values in baru seedlings of 26.4 g per kg, which falls within this range. While evaluating Baru’s nutritional status, Souza et al. (2018) found that suitable N levels for seedling development ranged from 20.20 to 26.00 g per kg.

**Table 2.**
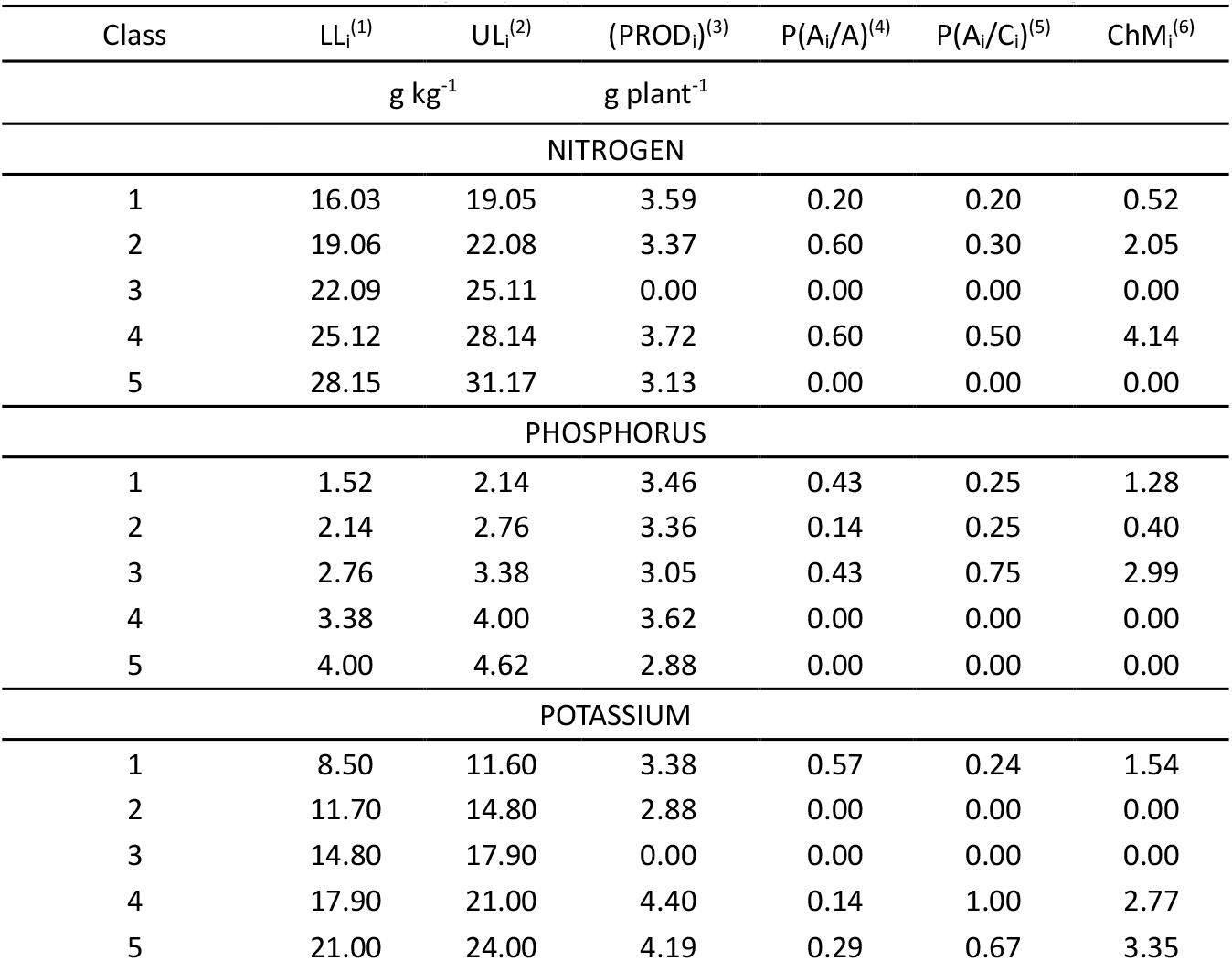
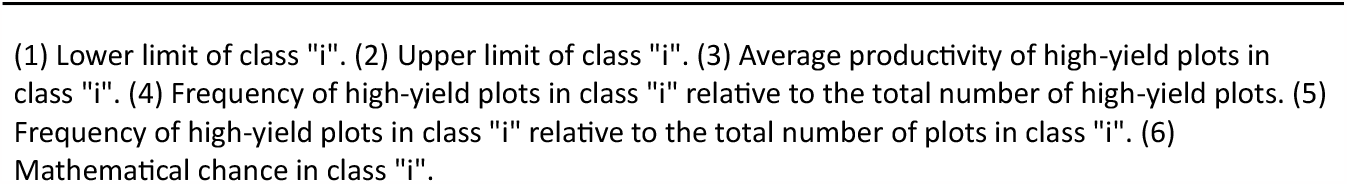
Reference values for nitrogen, phosphorus, and potassium for baru seedlings.

For the determination of the sufficiency range of phosphorus (P), adopted a similar criterion as for N. Thus, grouped the range of classes 1 to 3, resulting in a sufficiency range of 1.52 to 3.38 g of P per kg (Table 2), which is similar to the values observed by Boni et al. (2016) and Souza et al. (2018).

Potassium (K) was grouped based on ChM values greater than 2.77 g per plant, encompassing classes 4 and 5, with sufficiency limits ranging from 17.90 to 24.00 g of K per kg (Table 2). The sufficiency range values were higher than the content of 10.8 g per kg obtained by Boni et al. (2016) and lower than the 28.04 g per kg reported by Souza et al. (2018). The range determined for K by ChM differed from the values reported in previous studies on baru nutrition, highlighting the need to establish a reference for seedling production of this species.

Calcium (Ca), magnesium (Mg), and sulfur (S) were nutrients that obtained narrow sufficiency range selections based on the higher ChM values. However, similar to the approach taken for nitrogen (N), a broader grouping of classes for Mg and S. For Mg, the sufficiency range covered classes with ChM values greater than 0.30 g per plant, and for S, ChM values greater than 0.70 g per plant. Only for Ca, it was not possible to extend the sufficiency range amplitude (Table 3).

**Table 3.** Reference values for calcium, magnesium, and sulfur for baru seedlings.

| Classe | LL<sub>i</sub><sup>(1)</sup> | UL<sub>i</sub><sup>(2)</sup> | (PROD<sub>i</sub>)<sup>(3)</sup> | P(A<sub>i</sub>/A)<sup>(4)</sup> | P(A<sub>i</sub>/C<sub>i</sub>)<sup>(5)</sup> | ChM<sub>i</sub><sup>(6)</sup> |
| --- | --- | --- | --- | --- | --- | --- |
|  | g kg<sup>-1</sup> |  | g plant<sup>-1</sup> |  |  |  |
| CÁLCIO |  |  |  |  |  |  |
| 1 | 1.76 | 3.54 | 3.56 | 0.57 | 0.33 | 2.41 |
| 2 | 3.54 | 5.32 | 2.97 | 0.00 | 0.00 | 0.00 |
| 3 | 5.32 | 7.10 | 4.52 | 0.29 | 0.00 | 0.00 |
| 4 | 7.10 | 8.88 | 3.28 | 0.00 | 0.00 | 0.00 |
| 5 | 8.88 | 10.66 | 3.57 | 0.14 | 0.50 | 0.91 |
| MAGNÉSIO |  |  |  |  |  |  |
| 1 | 0.74 | 1.12 | 3.43 | 0.57 | 0.33 | 2.24 |
| 2 | 1.12 | 1.50 | 3.54 | 0.14 | 0.17 | 0.30 |
| 3 | 1.50 | 1.88 | 3.86 | 0.29 | 0.00 | 0.00 |
| 4 | 1.88 | 2.26 | 3.97 | 0.00 | 0.00 | 0.00 |
| 5 | 2.26 | 2.64 | 3.02 | 0.00 | 0.00 | 0.00 |
| ENXOFRE |  |  |  |  |  |  |
| 1 | 1.26 | 1.76 | 3.43 | 0.57 | 0.31 | 2.07 |
| 2 | 1.76 | 2.26 | 3.83 | 0.14 | 0.33 | 0.70 |
| 3 | 2.26 | 2.76 | 3.65 | 0.14 | 0.00 | 0.00 |
| 4 | 2.76 | 3.26 | 2.45 | 0.00 | 0.00 | 0.00 |
| 5 | 3.26 | 3.76 | 4.17 | 0.14 | 0.50 | 1.24 |

(1) Lower limit of class "i". (2) Upper limit of class "i". (3) Average productivity of high-yield plots in class "i". (4) Frequency of high-yield plots in class "i" relative to the total number of high-yield plots. (5) Frequency of high-yield plots in class "i" relative to the total number of plots in class "i". (6) Mathematical chance in class "i".

The selection of the appropriate range for Ca was at 2.41 g per plant, and it was not feasible to group classes for Ca, resulting in a narrow sufficiency range with a low amplitude between the minimum (1.76 g per kg) and maximum (3.54 g per kg) leaf content. When comparing these results with those of Souza et al. (2018), the obtained Ca values were much lower, ranging from 22.39 to 24.77 g per kg in their study. Boni et al. (2016) and Silva et al. (2019) reported nutritional levels consistent with the sufficiency range values obtained by ChM.

The sufficiency ranges established for magnesium (Mg) were 0.74 to 1.50 g per kg, covering classes 1 and 2, and similarly, for sulfur (S), the classes determined by ChM were 1 and 2, with values of 1.26 to 2.26 g per kg. These values are similar to the leaf content levels found for baru by Boni et al. (2016), Silva et al. (2019), and Souza et al. (2018), confirming the optimal leaf nutrient levels for the crop.

In the same way that ChM values for macronutrients, the chance of leaf nutrient classes was estimated for micronutrients in baru seedlings (Table 4).

**Table 4.**
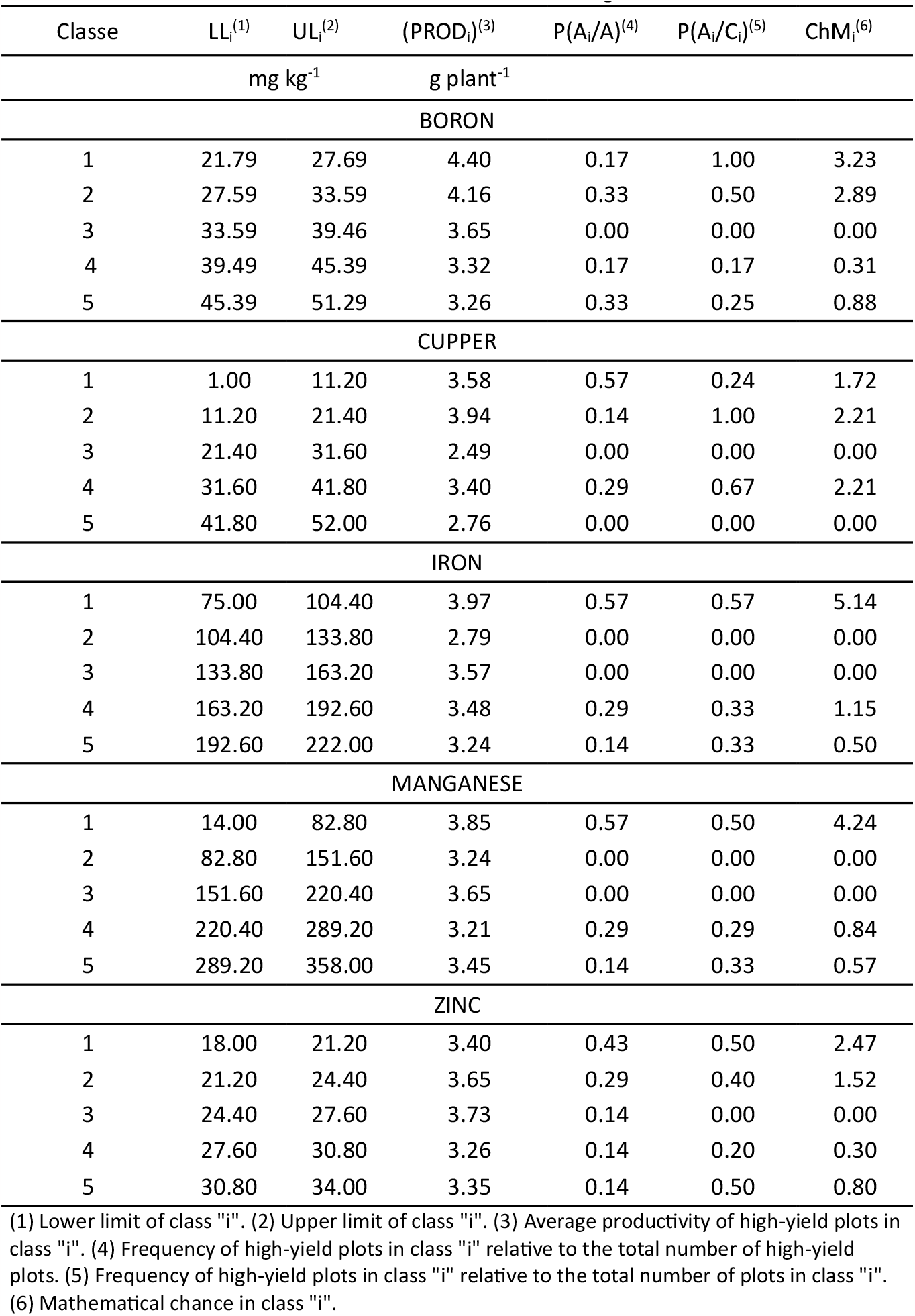
Reference values for micronutrients for baru seedlings.

Boron (B) was grouped based on ChM values greater than 2.89 g per plant, encompassing classes 1 and 2, resulting in sufficiency range limits of 21.79 to 33.59 mg of B per kg (Table 4).

These values were similar to the 14.7 to 33.0 mg per kg reported by Boni et al. (2016) and lower than the 49.38 mg per kg reported by Souza et al. (2018).

The optimal range for copper (Cu), obtained through the mathematical chance method, involved grouping classes 1 and 2 with a sufficiency range of 1.0 to 21.4 mg per kg. The amplitude of this range was higher than the values reported by Boni et al. (2016) and Souza et al. (2018). However, the copper nutritional levels obtained in these studies fell within the range determined by ChM. The wide range for Cu suggests that baru has a low requirement for this nutrient for production.

The values for the optimal range of iron (Fe) and manganese (Mn) tended to result in a narrow range. In both cases, only one class (Class 1) was selected based on ChM values; however, more than half of the observations with high MSPA production were in these groups (Table 4). For Fe, the optimal range had limits of 75.0 to 104.4 mg per kg, with values reported by Boni et al. (2016) and Souza et al. (2018) exceeding the sufficiency range obtained by ChM. In the natural conditions of the Cerrado soils, Fe is not a limiting factor for growth. Still, determining sufficiency ranges can assist in fertilization planning for seedling production.

Manganese (Mn) had a sufficiency range with limits of 14.0 to 82.8 mg per kg, with values reported by Souza et al. (2018) and Boni et al. (2016) exceeding 200 mg per kg, well above the upper limit of the range determined by ChM.

The values for the optimal range of zinc (Zn) covered classes 1 and 2, with optimal levels from 18.0 to 24.40 mg per kg, which were higher than the reference levels in studies on baru (SOUZA et al., 2018; BONI et al., 2016). Determining the range for Zn included a large proportion of samples, as more than 70% of the samples with high MSPA production fell within this range.

In general, the sufficiency ranges obtained through the ChM method for baru seedlings were close to values found in the literature. However, using the ChM method allowed for assessing and establishing sufficiency ranges for the crop.

## Conclusions

The sufficiency ranges determined using the ChM method for macronutrients were as follows in g per kg:

Nitrogen (N): 19.06 to 28.14

Phosphorus (P): 1.52 to 3.38

Potassium (K): 17.9 to 24.0

Calcium (Ca): 1.76 to 3.54

Magnesium (Mg): 0.74 to 1.50

Sulfur (S): 1.26 to 2.26

For micronutrients, the sufficiency ranges obtained through the ChM method were as follows in mg per kg:

Boron (B): 21.79 to 33.59

Copper (Cu): 1.0 to 21.40

Iron (Fe): 75 to 104.4

Manganese (Mn): 14.0 to 82.8

Zinc (Zn): 18.0 to 24.4

